# Anxiolytic activity and phytochemical content of the methanol extract of *Blumea balsamifera* (L.) DC. in ICR mice

**DOI:** 10.64898/2026.07.25.740683

**Authors:** Jazzy Jane Alberca, Krisha Fe N. Limosnero

## Abstract

Anxiety disorders significantly impact quality of life worldwide. While anxiolytic drugs like benzodiazepines are effective, they pose risk of adverse effects. This has driven interest in plant-derived alternatives with safer profiles. *Blumea balsamifera* (L.) DC. is traditionally used in Southeast Asia for medicinal purposes, yet its anxiolytic activity has not been scientifically evaluated. This study investigated the anxiolytic potential of the methanol extract of *B. balsamifera* leaves in male Institute of Cancer Research mice using the elevated plus maze test. This was an experimental, oral dose-response study employing a randomized controlled design to compare the effects of *B. balsamifera* against control groups. The extract underwent photochemical screening and an acute oral toxicity test following OECD Guideline 423. Subsequently, anxiolytic activity was evaluated using the elevated plus maze test at oral doses of 250, 500, or 1000 mg/kg. Diazepam (1 mg/kg) and 1% Tween 80 (10 mL/kg) served as the positive and negative control, respectively. A power analysis determined a minimum sample size of 25 animals (five groups) to achieve ≥ 0.95 power (α = 0.05; f = 1). Phytochemical screening identified the presence of flavonoids, tannins, terpenoids, and phenols. Acute oral toxicity test indicated an estimated LD₅₀ of 5000 mg/kg. In the elevated plus maze test, ICR mice treated with 500 and 1000 mg/kg showed a statistically significant increase compared to the negative control. These findings demonstrate a favorable safety profile and dose-dependent anxiolytic-like effects, providing the like effects, providing the first scientific evidence supporting *B. balsamifera* leaves as a potential natural anxiolytic agent.

## Introduction

Many individuals experience occasional anxiety, such as worrying about health, family, or career. However, when anxiety becomes persistent and overwhelming, it can develop into a disorder, affecting both mental and physical well-being [1]. Anxiety disorders were reported to affect approximately 4% of the global population, making them the leading mental, emotional, and behavioral issues, with a rising incidence observed among the youth. In the Philippines, more than six million individuals were reported to have experienced anxiety and depression [2], highlighting the pressing need for accessible and effective treatment options [3].

Anti-anxiety medications evolved from early sedatives like barbiturates and meprobamate to safer, more targeted treatments such as benzodiazepines, and selective serotonin or norepinephrine reuptake inhibitors (SSRIs and SNRIs). As understanding of the neurobiology of anxiety deepened, newer multi-target compounds were developed [4,5]. Nevertheless, many anxiolytic drugs still posed risks including side effects, delayed onset, and dependency risks. This led to growing interest in plant-based alternatives with fewer adverse effects, which often contain central nervous system-active compounds [6]. Broadening the scientific study of these plants strengthens the potential to discover effective natural remedies.

Validation of plant-based treatments was conducted through animal models. While animal models do not reproduce the full range of symptoms seen in human anxiety disorders, they are useful for inducing a general anxious state for scientific study [7]. Despite limitations, these models remain an indispensable tool for probing the neurobiological foundations of anxiety and testing potential therapeutic interventions in controlled, reproducible settings. The elevated plus maze (EPM) remained one of the most widely used and pharmacologically validated model for screening anxiolytic activity due to its simplicity, reproducibility, and sensitivity [7,8]. Designed to capture rodents’ natural conflict between exploring novel environments and avoiding open and elevated areas, EPM behavior reliably reflects anxiety levels [8,9]. This model consistently responded to standard anxiolytics targeting GABA and glutamate systems in the brain [7,10].

Rodents, especially mice and rats, are widely used in anxiety research because of their genetic, neuroanatomical, and behavioral similarities to humans. They are favored for their well-studied nervous systems, ease of handling, and cost-effectiveness. While many anxiety models were originally developed in rats, they have since been adapted for mice, with varying degrees of success. Mice are particularly valuable in genetic studies due to their susceptibility to genetic modification [11]. A notable limitation in preclinical research is the strong sex bias, with male rodents used over ten times more frequently than females. This preference stems partly from the hormonal cycles in females, which can introduce variability in experimental results. Despite the higher prevalence of anxiety disorders in women, this male bias continues due to concerns about the added complexity of female hormonal fluctuations and differences in drug metabolism and response [9,12]. Institute of Cancer Research (ICR) mice were selected for this study because they represent an established and well-validated outbred model for evaluating neuropsychiatric phenotypes and screening anti-anxiety interventions [13].

Several plants have been studied using EPM and similar tests. *Blumea lacera* (Burm.f.) DC. exhibited anxiolytic effects in the hole board test, open field test, elevated plus maze, and light and dark box test [14,15]. Similarly, *Blumea densiflora* DC. showed anxiolytic effects in the hole cross test, open field test, and swing test [16]. These findings suggest that members of the *Blumea* genus may contain phytochemicals capable of reducing anxiety.

One species of interest is *Blumea balsamifera* (L.) DC., locally known as “sambong” in Tagalog or “gabon” in Cebuano. This plant is widely grown across the Philippines and has long been used in traditional medicine to treat a range of conditions, including infections, inflammation, kidney stones, and liver disorders [3,17]. Its therapeutic properties have been attributed to its diverse bioactive components. However, despite its known pharmacological value, the anxiolytic potential of *B. balsamifera* has not yet been scientifically investigated.

This study aimed to determine the anxiolytic potential of the crude methanol extracts of *B. balsamifera* in Institute Cancer Research (ICR) strain male albino mice. Specifically, the crude methanol extract of the leaves of *B. balsamifera* was phytochemically screened, determined for its non-lethal concentration by performing an acute oral toxicity test, and its anxiolytic effects were compared with the standard anxiolytic drug, diazepam. This study offers valuable contributions across several fields. In pharmacognosy, the study contributes to the ongoing search for bioactive compounds in medicinal plants by adding to the growing body of knowledge on plant-based therapies, particularly targeting anxiety-related conditions. In pharmaceutical sciences, this study aligns with current efforts to find safer, more accessible, and cost-effective alternatives to conventional anxiolytic medications. By investigating *B. balsamifera*’s potential anxiolytic effects, this study provides an empirical, behavioral foundation for traditional medicine and modern mental health research. This furthermore supports the local communities through the sustainable and informed use of a native plant with known medicinal and cultural significance. Through this research, baseline data on *B. balsamifera*’s anxiolytic activity may be established that may support the development of accessible, plant-based treatments for anxiety.

## Methodology

### Sample collection

Five kilograms of mature and healthy leaves of *Blumea balsamifera* (L.) DC., locally known as “Sambong” in Tagalog or “Gabon” in Bisaya, were collected from Guso, Jomgao, Argao, Cebu, Philippines, at coordinates of 9°53’54.7”N and 123°34’49.8”E. The exact sampling location is illustrated in Fig 1a, while Fig 1b presents the *B. balsamifera* plant at the sampling site. The identity of the plant was authenticated by Mr. Val B. Salares from the Department of Biology at the University of San Carlos (see S1 Appendix).

**Fig 1.**
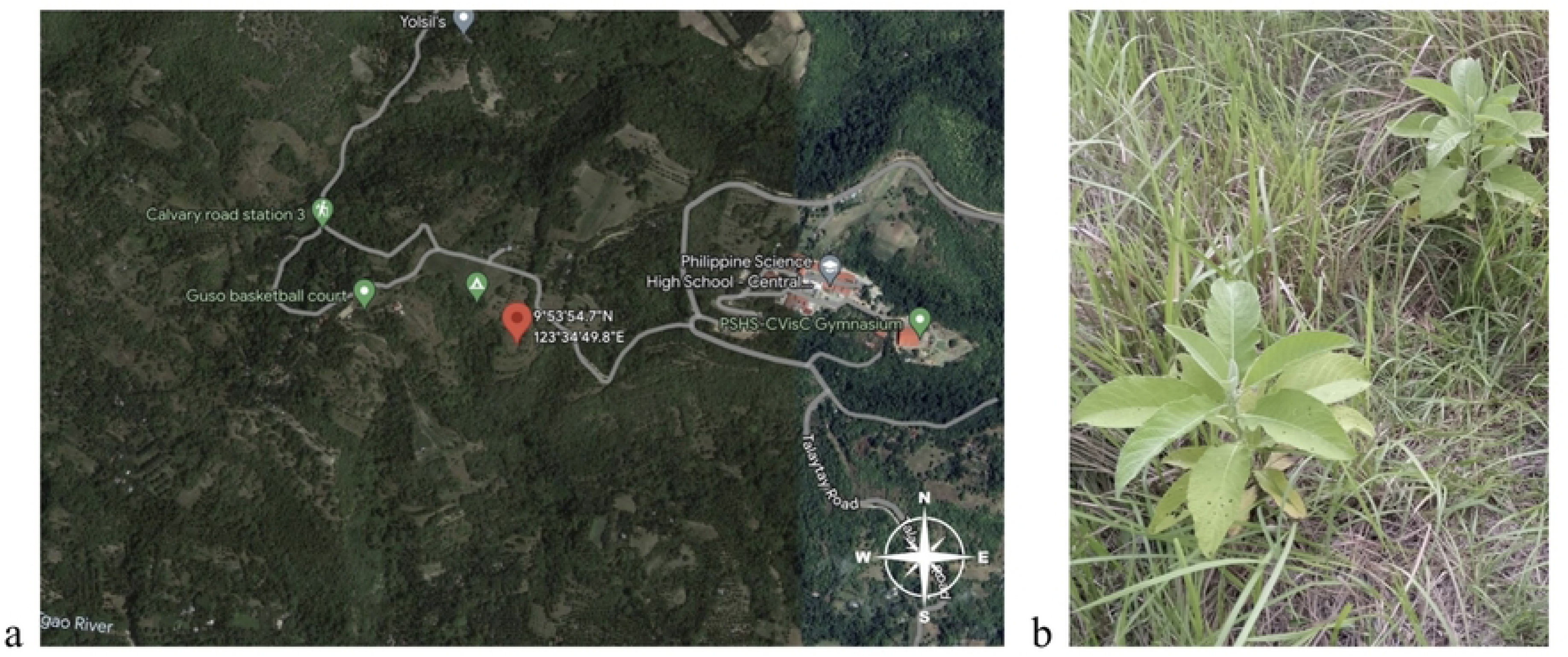
(a) Sampling location in Guso, Jomgao, Argao, Cebu, Philippines; (b) *B. balsamifera* at sampling site.

### Sample preparation

Freshly collected *Blumea balsamifera* (L.) DC. were washed with distilled water to remove adhering dirt and impurities. The leaves were then air-dried at room temperature for approximately 8–10 days, following the procedures of Jirakitticharoen *et al.* [18] and Sakee *et al.* [19]. The dried leaves were ground into a coarse powder using a blender. The resulting powder was packed in ziplock bags and analyzed for moisture content with a moisture analyzer. In this study, the moisture content was monitored, and drying was continued in a desiccator until moisture content dropped below 10%, the powder was transferred from the ziplock bags and stored in the desiccator until extraction. Measuring the moisture content helps ensure that the extract yield is based on the actual dry weight of the plant material, allowing for more accurate, consistent, and comparable results throughout the study. Sample preparation followed the method described by Hossain *et al.* [16], with adaptations specific to this study.

### Extraction process

The extract was prepared by soaking 100 grams of powdered, dried *Blumea balsamifera* (L.) DC. leaves in 1000 mL of 100% methanol for 48 hours. The sample-to-solvent ratio used in this study was 1:10, following Toralba *et al.* [20]. The filtrate was dried at 40°C under vacuum using a rotary evaporator and further concentrated by evaporation. This extraction process was repeated a total of three times using the same plant material and methanol used in the first extraction to maximize the yield of the crude extract. The resulting crude methanol extract contained a clear yellowish oil and a green gummy residue. Some of the oil formed a distinct layer that was easily separated by decantation, while the remaining oil emulsified with the residue could not be removed by this method. To facilitate further separation, the extract was stored at 4°C, which reduced the solubility of the emulsified oil and caused it to aggregate into visible droplets or clumps. The aggregated oil was skimmed off, and the extract was stored again at 4°C until use in the experiment [14].

### Phytochemical screening

The crude methanol extract of *Blumea balsamifera* (L.) DC. leaves was analyzed for the presence of alkaloids, flavonoids, steroids, triterpenoids, terpenoids, tannins, saponins, and phenols using the standard methods described by Shaikh and Patil [21] and Doughari and Saa-Aondo [22] (see S2 Appendix). The positive controls for each test were the commonly used reference compounds, while distilled water served as the negative control for all tests.

### Experimental animals

All procedures involving mice were conducted at the Pharmacology Laboratory of the University of San Carlos (USC) Department of Pharmacy, and any procedures performed or incidents that occurred during the experiment were reported to or discussed with the USC Laboratory Animal Facility (LAF) technician. Approval from the USC Institutional Animal Care and Use Committee (Protocol Number: 2024–056–028) was obtained prior to conducting any biological studies (see S3 Appendix).

#### Animal acquisition

A total of thirty-seven (37) male albino mice of the Institute of Cancer Research (ICR) strain, aged six to eight weeks and weighing between 15 and 29 g, were acquired from the USC LAF. Of these, twenty-five (25) mice were allocated to the elevated plus maze test. Mice were randomly assigned to their respective groups by the USC LAF technician using a simple randomization method, such as online computer-generated random numbers or drawing lots, to ensure each animal had an equal chance of being allocated to any group. The mice were then placed into five (5) cages, with five (5) mice housed per cage. The principal investigator then randomly assigned these cages to the following treatment groups: a negative control, positive control, and three (3) test groups. The remaining twelve (12) mice were allocated to the acute oral toxicity test and were randomly grouped, using the same randomisation method, into four (4) separate cages, with three (3) mice housed per cage. Each individual mouse across the entire study was assigned an identifier, such as a number written on its tail.

#### Acclimatization and handling

All experimental procedures were conducted following a 7-day acclimatization period. Proper personal protective equipment, including laboratory gown, hairnet, mask, and gloves, was required inside the pharmacology laboratory. The working area was cleaned both during and after use. All individuals interacting with the mice thoroughly washed their hands upon entering and leaving the room.

#### Euthanasia

At the end of the study, euthanasia was performed using carbon dioxide inhalation and cervical dislocation. The carcasses were then autoclaved and temporarily stored in a freezer prior to incineration.

### Animal housing and husbandry

This procedure was adapted from Walf and Frye [23] and complied with the USC Laboratory Animal Facility guidelines. The mice were housed under conventional housing conditions at the Animal House of the Department of Pharmacy, University of San Carlos, with access restricted to authorized personnel. Mice were group-housed in polycarbonate cages (18 cm × 10 cm × 7 cm) lined with autoclaved wood shavings as bedding. Five (5) cages were used for the elevated plus maze test (5 mice/cage), while four (4) cages housed the mice allocated to the acute oral toxicity test (3 mice/cage). Throughout the experiment, mice were maintained under a 12-hour light/dark cycle with the light phase starting at 7:30 AM. The temperature was controlled at 21 ± 3°C and relative humidity was maintained between 40% and 70%. Each mouse was provided daily with 5-10 g of a standard laboratory rodent pellet diet (18-20% crude protein, 4-5% fat, 4-6% fiber; SmartHeart Rodent Pellets) and clean, potable drinking water *ad libitum*. Environmental enrichment—including nesting material, paper tubes, or shelters—was provided to promote normal exploratory and nesting behaviors. Routine sanitation procedures were followed; cages were cleaned, water bottles were refilled with fresh water, and both bedding and nesting materials were replaced twice weekly.

### Treatment administration

#### Dosage preparation

Stock solution preparations and dose volume followed the guidelines of Erhirhie and Ajaghaku [24] and Pandy [25]. The stock solution was prepared at one-tenth of the desired dosage (S4 Appendix). The administered volume did not exceed 20 mL/kg b.w., as specified by the Organisation for Economic Co-operation and Development [26] guidelines, for aqueous solvents. A constant volume was maintained across dosages by adjusting the conc. of the stock solution (S5 Appendix).

#### Oral gavage procedure

The prepared doses of the crude methanol extract of *Blumea balsamifera* (L.) DC. were administered orally (p.o.) as a single dose via oral gavage using a feeding tube. Mice were manually restrained to minimize stress and to ensure accurate dosing. This was achieved by gently holding the mouse at the base of the tail and lifting it. One hand secured the mouse’s body by cupping it in the palm, while the other hand held the mouse’s head steady before insertion of the gavage tube.

### Acute oral toxicity test

The protocol described in the Organisation for Economic Co-operation and Development (OECD) Guideline 423 was followed [26]. This method classifies the substances into toxicity categories based on fixed LD_50_ cut-off values and requires at least two doses showing mortality between 0% and 100%. While it does not provide a precise LD_50_ value, the test identifies exposure ranges where lethality is expected, as mortality is the primary endpoint.

#### Number of animals and dose levels

The randomly allocated male mice (n = 3) were administered a single oral dose of the crude methanol extract of *Blumea balsamifera* (L.) DC in a stepwise manner at dosages of 5, 50, 300, and 2000 mg/kg body weight, starting with 300 mg/kg body weight due to the lack of prior information on the toxicity of the methanol extract of *B. balsamifera*. Mortality at each step guided the next dose, leading to either discontinuation of further testing, repetition of the dose, or adjustment to a higher or lower level. The test scheme began with a dose of 300 mg/kg body weight. The time interval between dosing groups depended on the onset, duration, and severity of toxic signs. Treatment at the next dose level was delayed until the survival of previously dosed animals were confirmed.

#### Dose administration

Mice were fasted for 3–4 hours before dosing, with food withheld for 1–2 hours post-dosing. Each mouse was monitored individually during the first 30 minutes after dosing, periodically over the first 24 hours (especially in the first 4 hours), and daily for 14 days.

#### Monitoring and observations

Daily monitoring and observations included assessments of the mouse’s general clinical condition such as changes in movement, coat condition, posture, breathing, feces and urine, attitude, appearance, and body condition scores (BCS). BCS was assessed through visual inspection and gentle palpation of the spine and hips to evaluate fat and muscle coverage, providing a quick indication of the mouse’s overall health and nutritional status. A 1-5 scoring scale was used, where 1 indicated an emaciated state, 3 represented ideal body condition, and 5 indicated obesity. The BCS scoring criteria and reference images are provided in S6 Appendix. Specific clinical signs such as tremors, convulsions, salivation, diarrhea, lethargy, and coma were also monitored. The monitoring and observation period was extended if toxic effects were delayed or if recovery took longer than expected. The timing of toxic signs, including their onset and resolution, was systematically documented. In accordance with the Humane Endpoints Guidance Document [26], animals in a moribund condition or experiencing severe pain or distress were humanely euthanized, and the time of death was precisely noted. Body weights were measured before dosing and daily for 14 days, with all changes documented. Detailed records of general clinical condition, specific clinical signs, weight changes, and mortality were maintained.

#### Gross necropsy

Although mortality is the primary endpoint of OECD Guideline 423, gross necropsy provides supplementary data on potential subclinical effects of the test substance. At the end of the observation period, all surviving mice were weighed, humanely euthanized, and dissected for gross necropsy to detect any subtle or hidden toxic effects not evident during daily monitoring and observations. The procedure was carried out with the assistance of a licensed veterinarian, Dr. Maxfrancis G. Talle, to ensure accurate identification and interpretation of pathological findings. Prior to dissection, the overall condition of each mouse, including BCS and musculature, was assessed. Vital organs such as the liver, kidneys, spleen, intestine, and heart were examined macroscopically for any deviations in color (e.g., pallor, darkening, mottling), size (e.g., swelling, atrophy, asymmetry), or texture (e.g., firmness, friability, nodules). Under normal conditions, these organs appear smooth and moist, with uniform color and consistency, specifically, the liver should be reddish-brown, the kidneys and heart firm and pink, the spleen dark red, and the intestines pale pink without distention or discoloration. Any irregularities were noted as potential indicators of inflammation, necrosis, hemorrhage, or other pathological changes. In addition, the organ weights of the liver, kidneys, and spleen were recorded.

### Elevated plus maze test

#### Treatment groups

The mice were randomly divided into groups, each containing five replicate mice (n = 5): the negative control, positive control, and test group. The negative control group received the vehicle, a 1% Tween 80 water solution (10 mL/kg b.w., p.o.). The positive control group was administered with the anxiolytic drug, diazepam (1 mg/kg b.w., p.o.). Diazepam was obtained from the University of Cebu Medical Center (UCMed) and was used with authorization certified by Dr. Rose May D. Camalongay of Camalongay Clinic (see S7 Appendix). The test group was further divided into three (3) subgroups, each receiving a different dosage of the crude methanol extract of *Blumea balsamifera* (L.) DC. The selected dosages were 250, 500, and 1000 mg/kg b.w., based on the results of the acute oral toxicity test. All dosed mice in the study had no prior exposure to the EPM test.

#### Setup and conditions

The elevated plus maze (EPM) apparatus, constructed from white melamine-faced birch plywood, consisted of two open arms (30 × 5 cm) and two closed arms (30 × 5 × 15.25 cm) with an open roof, and was elevated 40 cm above the floor. The maze was placed in the Pharmacology Laboratory, where the mice were tested individually. The “Timestamp Camera Basic” app on a smartphone was used as a video recorder and timer, positioned above the maze. Natural light with room lights turned off was maintained throughout the experiment for consistency. Plastic tables were stacked on their sides around the EPM apparatus. These tables also served as the physical barrier, separating the experimenter from the testing area and minimizing the likelihood that the mice would react to human presence. The testing period was scheduled between 1:00 PM and 5:00 PM to align with the light phase of the mice’s circadian rhythm when anxiety-like behaviors are more stable and reproducible [8,27].

#### Behavioral testing procedure

Thirty (30) minutes after oral gavage administration, each mouse was removed from its cage and placed at the center of the maze, facing the open arm opposite the experimenter. After placing each mouse in the center of the maze, the experimenter moved quietly behind the barrier and remained still for the duration of the 5-minute recording. The timer was started when the mouse was placed in the maze and was set for five (5) minutes. At the end of the 5-minute test, the mouse was removed from the maze and returned to its cage. The apparatus was cleaned with 70% ethanol and dried with paper towels to eliminate any olfactory cues before testing the next mouse. To maintain consistency in maze exposure, mice that fell off the maze or froze for prolonged periods in the open arms were excluded from the analysis, as these behaviors indicate heightened anxiety and could compromise data integrity. The procedure was based on Walf and Frye [23].

#### Behavioral parameters

The following parameters were documented during the 5-minute recording: (i) the number of entries into the open and closed arms, and (ii) the average time spent in the open and closed arms. An entry was defined as all four (4) paws being within an arm. To account for variations in general activity, the percentage of open arm time (%OAT) and open arm entries (%OAE) were calculated as the ratio of open arm activity to total arm activity. These activities were compared across groups. Every precaution was taken to ensure that external stimuli, e.g., noise, bright illumination, or visible human presence, were minimized to avoid inducing anxiety in the mice.

### Statistical Analysis

A power analysis for a One-way ANOVA (fixed effects, omnibus) was performed using G*Power, shown in Fig 2 [28]. The power analysis indicated that a minimum sample size of 25, distributed across five (5) groups, was required to achieve a statistical power of at least 0.95 with an alpha of 0.05 and a large effect size (f = 1). All experimental data was expressed as mean ± SEM and significance was established at *p* < 0.05. Statistically significant differences between groups were determined using a One-way analysis of variance (ANOVA), followed by post hoc Tukey’s multiple range tests [23], performed using SPSS.

**Fig 2.**
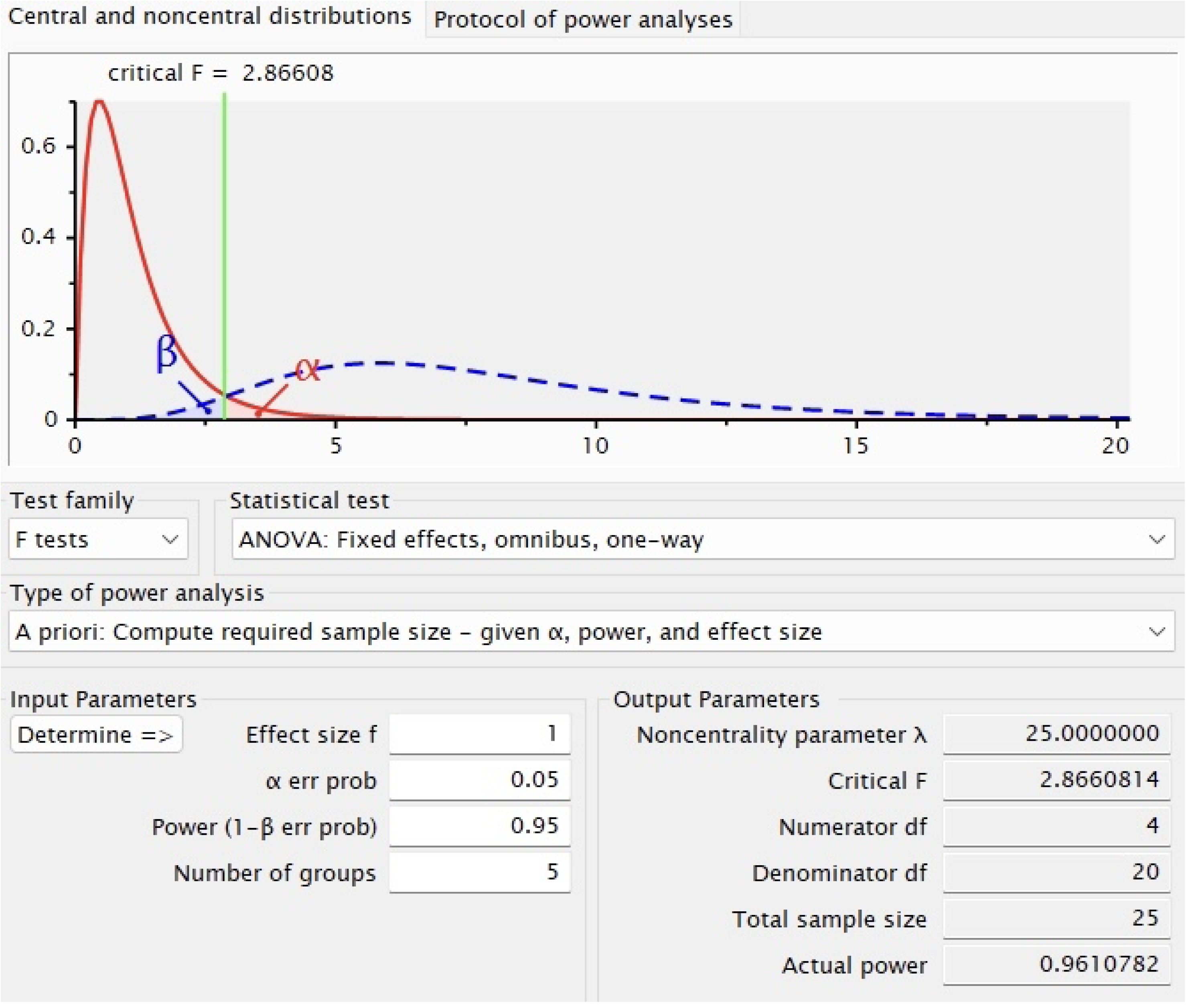
Power analysis for one-way ANOVA using G*Power.

## Results and discussion

### Preparation of crude methanol extract

From the air-dried *Blumea balsamifera* (L.) DC. leaves collected, a total of 128 grams of coarse leaf powder was obtained. The moisture content of the powder was determined to be 9.62% ± 0.15. The 128 grams of coarse leaf powder were soaked in 1.28 L 100% methanol for 48 hours. The resulting filtrate was concentrated at 40°C under reduced pressure using a rotary evaporator, followed by further evaporation. The combined crude methanol extract from three extraction cycles, excluding the oily fraction, yielded approximately 16 grams and was stored at 4°C until use. The presence of an oily fraction suggests the extraction of lipophilic constituents such as volatile oils, which are known to be abundant in *B. balsamifera* [29].

### Phytochemical screening

A qualitative phytochemical screening was performed on the crude methanol extract of *Blumea balsamifera* (L.) DC. leaves, and the results are summarized in Table 1. The analysis indicated the presence of flavonoids, terpenoids, tannins, and phenols, while alkaloids, steroids, triterpenoids, and saponins were not detected. These compounds, particularly flavonoids, terpenoids, tannins, and phenolic compounds, are consistently reported in the scientific literature as naturally occurring constituents of *B. balsamifera* [30,31]. Volatile compounds such as terpenoids and phenolic acids make up the largest portion of the plant’s phytochemical content, while flavonoids are regarded as the major non-volatile constituents, along with sterols and sesquiterpene lactones [29,30]. These compounds have been identified in previous studies through phytochemical screening, as well as through advanced analytical techniques such as gas chromatography-mass spectrometry and high-performance liquid chromatography.

**Table 1.** Phytochemical analysis of the crude methanol extract of *B. balsamifera* leaves.

| Phytochemicals | Result |
| --- | --- |
| Alkaloids | - |
| Flavonoids | + |
| Steroids | - |
| Triterpenoids | - |
| Terpenoids | + |
| Tannins | + |
| Saponins | - |
| Phenols | + |

Signs indication: (+) = present, (-) = absent

In previous phytochemical screening studies, tannins have also been consistently detected when extracted using both polar solvents like methanol, ethanol and ethyl acetate and in non-polar solvents like n-hexane. Flavonoids, terpenoids, phenols, saponins, and steroids have most often been extracted using polar solvents, while phenols, steroids, triterpenoids, and alkaloids have also been reported in non-polar extracts [32–34]. The presence of flavonoids, terpenoids, tannins, and phenolic compounds in the current study supports their natural occurrence in *B. balsamifera*, as widely established in scientific literature. The absence of alkaloids, triterpenoids, saponins, and steroids in this extract may be attributed to their low abundance or variations in extractability, which depend on factors such as solvent used, plant part analyzed, or environmental conditions during growth.

Each of the phytochemicals detected has been widely studied for its roles in anxiety reduction through various biochemical mechanisms. Flavonoids and terpenoids demonstrated the ability to enhance GABAergic neurotransmission, similarly to benzodiazepines, by facilitating chloride ion influx and promoting neuronal inhibition [35]. Likewise, tannins and phenolic compounds have shown anxiolytic potential through their influence on both GABA and monoamine systems, which play critical roles in regulating mood and emotional responses [36].

Flavonoids, derived from polyphenolic structures, are known for their diverse biological actions, including effects on mental health. Numerous preclinical studies have explored their anxiolytic properties using behavioral models. These compounds exert their effects by interacting with various neurotransmitter pathways, enhancing neurotrophic activity, and reducing oxidative stress and inflammation within the central nervous system [37]. This group of compounds also possess antioxidant activity, which helped protect neural tissues and contributed to their anxiolytic effects [35]. The wide interest in flavonoids arose from the diversity in their chemical subtypes and the range of physiological systems they influenced [37].

Terpenoids, a broad class of secondary metabolites distributed across numerous plant families including Cannabaceae [38], had been reported to enhance cognitive functions, such as memory and learning, and to produce antidepressant and anxiolytic effects [39]. Many essential oils with known anxiolytic effects typically contained terpenoid alcohols like linalool, geraniol, and citronellol, as well as monoterpenes such as limonene or citral [40].

Tannins, classified as water-soluble polyphenols, were commonly found in various edible plants [41]. Although their anxiolytic effects had not been extensively studied in isolation, several investigations had demonstrated that extracts rich in condensed tannins produced anxiety-reducing effects through GABAergic mechanisms [42,43]. Additionally, hydrolysable tannins were shown to affect molecular pathways involved in stress and mood regulation [44].

Phenols constituted one of the most abundant and widely distributed classes of aromatic compounds in plants [45]. Flavonoids and phlorotannins, both types of phenolics, were recognized for their ability to modulate the benzodiazepine site of GABA_A_ receptors, contributing to sedative, anticonvulsant, and anxiolytic effects. Among these, flavonoids had drawn particular attention due to their therapeutic potential in managing anxiety and seizure-related conditions [46].

The identification of flavonoids, terpenoids, tannins, and phenols in the crude methanol extract of *B. balsamifera* leaves provide a strong chemical basis for the plant’s observed anxiolytic-like properties. Through various modes of action, including receptor modulation, antioxidant activity, and neurotransmitter regulation, these compounds collectively suggested that the *B. balsamifera* held promise as a natural source for anxiety management.

### Acute oral toxicity test

#### Number of animals and dose levels

The acute oral toxicity test was conducted according to the stepwise procedure [26]. Initially, three (3) mice were administered a dose of 300 mg/kg b.w., p.o. of the crude methanol extract of *Blumea balsamifera* (L.) DC. via oral gavage and observed for signs of toxicity and mortality over a 14-day period. As no mortality occurred, a second group of three (3) mice received the same dose. Based on the continued absence of mortality, the dose was escalated to 2000 mg/kg b.w., p.o., and administered to a third group of three (3) mice. Again, no mortality was recorded during the observation period. This led to a fourth group of three (3) mice being treated with the same dose, and no mortality was observed. Testing at a higher dose, such as 5000 mg/kg b.w., is typically unnecessary unless justified by a specific regulatory need, which refers to situations where national or international regulatory agencies require more precise hazard classification, risk assessment, or product registration data. For example, the U.S. Environmental Protection Agency may request data beyond the limit dose for chemicals intended for pesticide use or widespread environmental exposure. In the absence of such requirements, the 2000 mg/kg b.w. dose was deemed sufficient as the limit dose, fulfilling the criteria for assessing the acute oral toxicity of the extract under standard guidelines.

#### Monitoring and observations

Throughout the 14-day observation period across all four (4) groups, the two groups of mice were treated with 300 mg/kg b.w., p.o. exhibited normal general clinical condition, with no observable clinical signs and a steady weight gain. In contrast, the two groups treated with 2000 mg/kg b.w., p.o. also showed a normal general clinical condition with no observable clinical signs but experienced an initial decrease in body weight from day 0 (day of dosing) to day 7, followed by recovery and steady weight gain through day 14. S8 Appendix presents the individual observation sheets used to monitor general clinical condition, clinical signs of toxicity, and changes in body weight for each mouse from day 0 to day 14, while Table 2 provides a summary of these findings.

**Table 2.**
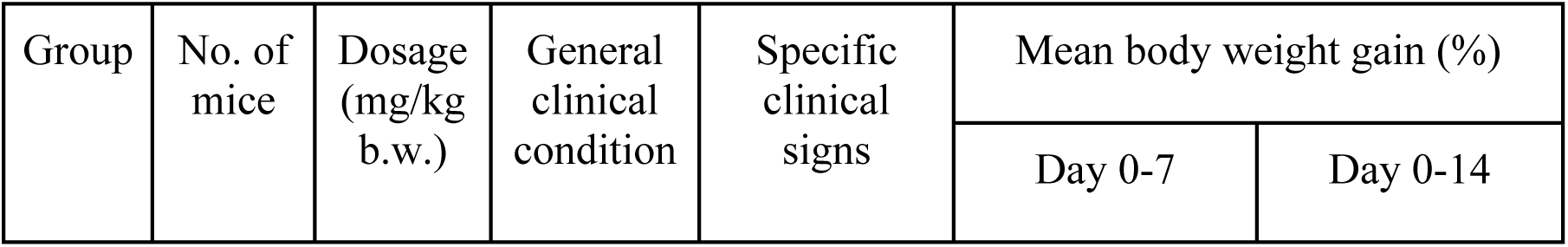
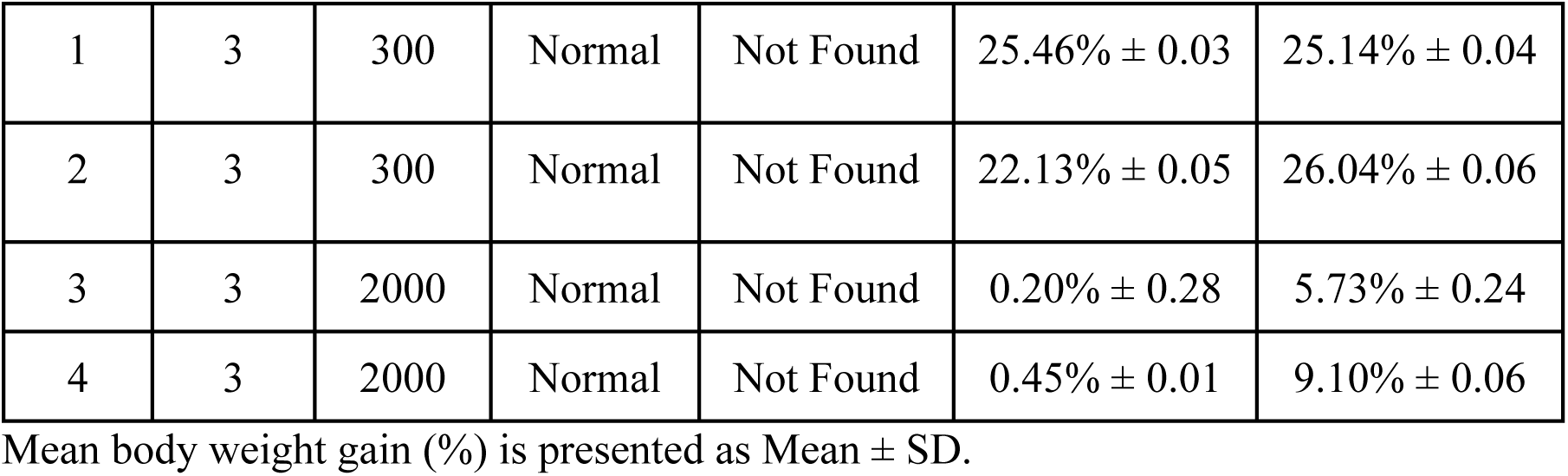
Summary of daily monitoring and observations of each mouse in each group treated with the crude methanol extract of *B. balsamifera* **leaves**.

| Group | No. of mice | Dosage (mg/kg b.w.) | General clinical condition | Specific clinical signs | Mean body weight gain (%) |  |
| --- | --- | --- | --- | --- | --- | --- |
|  |  |  |  |  | Day 0-7 | Day 0-14 |
| 1 | 3 | 300 | Normal | Not Found | 25.46% ± 0.03 | 25.14% ± 0.04 |
| 2 | 3 | 300 | Normal | Not Found | 22.13% ± 0.05 | 26.04% ± 0.06 |
| 3 | 3 | 2000 | Normal | Not Found | 0.20% ± 0.28 | 5.73% ± 0.24 |
| 4 | 3 | 2000 | Normal | Not Found | 0.45% ± 0.01 | 9.10% ± 0.06 |

Mean body weight gain (%) is presented as Mean ± SD.

Body weight gain is a general indicator of health, reflecting changes in muscle, fat, and fluid accumulation, as well as metabolic activity [47]. In acute oral toxicity studies involving mice, a weight loss exceeding 25% within 7 days is considered a sign of excessive toxicity or animal suffering. This threshold is one of the criteria used to evaluate toxicity, as described in OECD Guidance Document 19 and cited by van Berlo *et al*. [48]. In this study, that threshold was not exceeded in either the 300 mg/kg b.w., p.o. and 2000 mg/kg b.w., p.o. groups.

The high standard deviation in mean body weight gain observed in group 3 was due to two mice showing weight loss greater than 10% within 7 days, while one mouse exhibited unusually high weight gain—exceeding that observed in the 300 mg/kg b.w., p.o. groups. This variation may be linked to social dominance, as dominant mice tend to eat and drink more frequently and rest less than subordinate mice. Alternatively, the weight loss in some mice may indicate stress or limited food/water access, while the unusually high weight gain may reflect stress-induced overeating [49]. However, other contributing factors cannot be ruled out.

#### Gross necropsy

Gross necropsy was performed immediately after euthanasia. The overall condition of all mice prior to death and musculature were normal. Body condition scores were 3 which represents the ideal body condition. The appearance of vital organs such as the liver, kidneys, spleen, intestine, and heart appeared normal for all mice. Their appearance was consistent with previously documented findings of grossly normal presentations in acute oral toxicity studies by Jothy *et al.* [50] and Ito *et al.* [51], as well as confirmed by the veterinarian who assisted. Importantly, there were no signs of organ enlargement, surface indentations, or hemorrhagic lesions (see S9 Appendix).

#### Organ weights

In this study, the weights of the liver, kidneys, and spleen were found to be appropriate relative to the mice’s body size, as confirmed by the assisting veterinarian. Table 3 shows that the recorded organ weights fell within a range consistent with general reference values reported by Davies and Morris [52], whose data included mice of similar body weights to those used in this study. While the reference lacked detailed strain and age specifications, it served as a reasonable basis for comparison. Relative organ weights were not calculated, as the primary objective of this study was to determine the non-lethal concentration of the crude methanol extract of *B. balsamifera* leaves through acute oral toxicity test, rather than to assess detailed organ-specific toxicity. Nonetheless, it is widely recognized that significant deviations in body and organ weight can be early indicators of toxicity in the toxicological evaluation process [53]. The absence of marked differences in organ weights, alongside normal clinical observations and lack of mortality, suggests that the administered extract did not cause significant structural damage or pathological alterations in the vital organs examined.

**Table 3.**
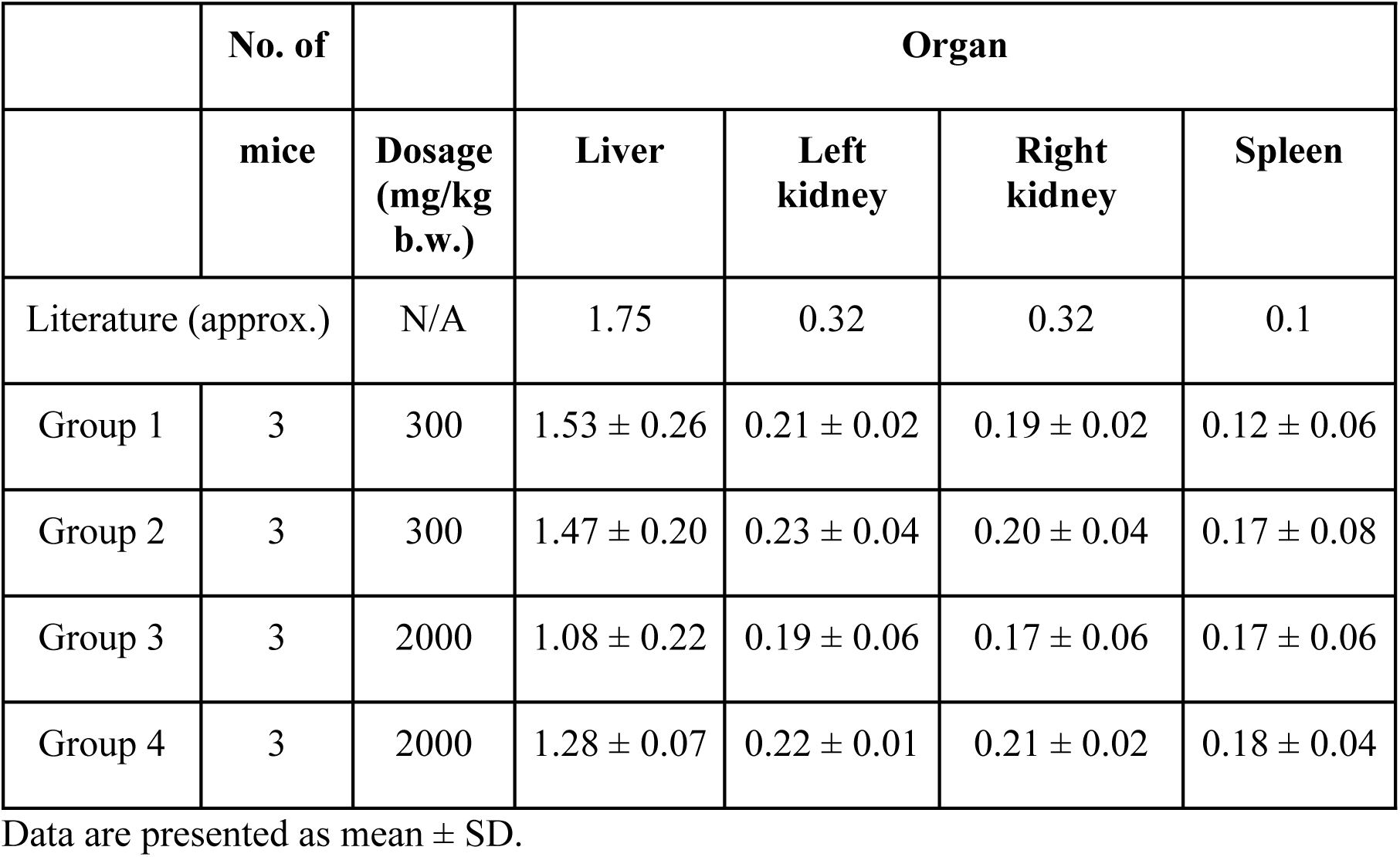
Organ weights of the vital organs examined during gross necropsy along with corresponding literature values for comparison.

Based on the test scheme [26], the crude methanol extract of *B. balsamifera* leaves was classified under Category 5 of the Globally Harmonized System for chemical classification, with an estimated LD₅₀ of 5000 mg/kg b.w. [26]. Following the dosing strategy used by Mesfin *et al.* [54], the estimated LD₅₀ value served as the basis for selecting doses for the elevated plus maze test. One-tenth of the estimated LD₅₀ was designated as the middle dose (500 mg/kg b.w.). From this, the low dose (250 mg/kg b.w.) and high dose (1000 mg/kg b.w.) were derived by halving and doubling the middle dose, respectively. These concentrations were used to evaluate anxiolytic activity of *B. balsamifera* in mice.

### Elevated plus maze test

Mice treated with 250, 500, and 1000 mg/kg b.w. of the crude methanol extract of *Blumea balsamifera* (L.) DC. (MeBB) leaves, administered orally (p.o.) via oral gavage, exhibited behavioral changes in the elevated plus maze (EPM) model that suggested a dose-dependent anxiolytic-like activity. Table 4 summarizes the behavioral responses at each dose compared with the negative control (10 mL/kg b.w., p.o., 1% Tween 80) and the positive control (1 mg/kg b.w., p.o., diazepam). All MeBB-treated and diazepam-treated groups showed increased average time spent in and number of entries into the open arms relative to the negative control, with statistically significant increases observed in MeBB_500_ and MeBB_1000_ groups. However, the MeBB_500_ group did not show a significant difference in the number of entries into the open arms. In terms of closed arms, the average time spent in all treatment groups showed a decreasing trend, with statistically significant reductions observed in the MeBB_500_, MeBB_1000_, and diazepam-treated groups relative to the negative control. The number of entries, on the other hand, increased across all treatment groups. These behavioral parameters supported the analysis of the percentage of open arm time spent (%OAT) and percentage of open arm entries (%OAE), which served as refined measures of anxiety-related behavior, adjusted for the mouse’s overall locomotor activity. Although testing conditions were standardized, some variability in behavioral responses was still observed. Converting raw data into percentage values helped minimize the effects of individual differences and general activity levels, allowing for more reliable comparisons across groups. However, residual variation likely resulted from uncontrolled biological factors such as individual health, body composition, gut microbiota, circadian rhythms, or genetic differences. Although behavioral scoring utilized high-precision video recording with millisecond timestamps, the principal investigator was not fully blinded to the treatment groups, introducing a potential source of observer expectation bias during split-second arm entry and exit decisions.

**Table 4.**
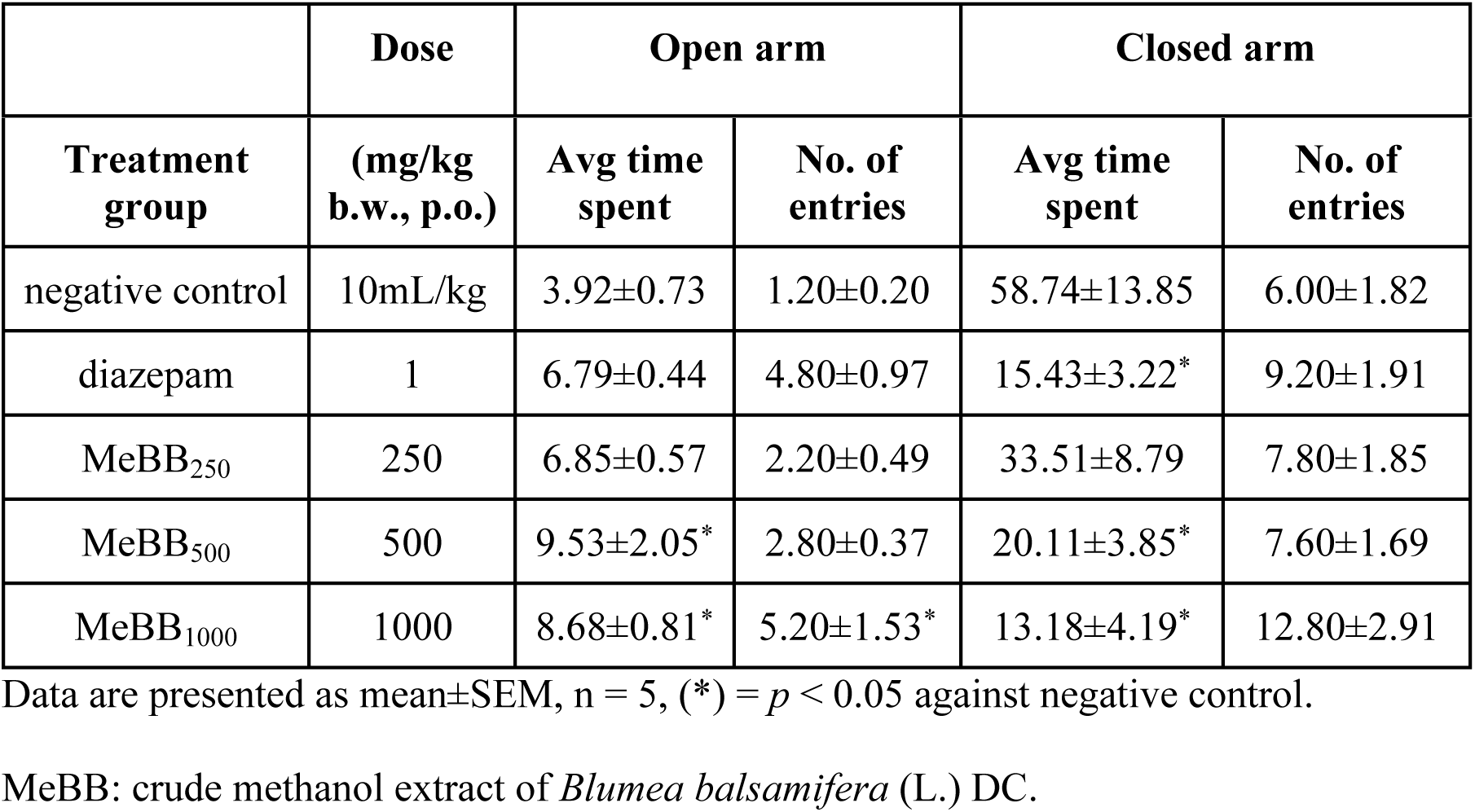
The behavior of mice treated with the crude methanol extract of *B. balsamifera* and reference compounds in the EPM model.

Fig 3 shows that the percentage of open arm time spent (%OAT) increased significantly in the MeBB_500_ (33.98% ± 7.15) and MeBB_1000_ (43.14% ± 4.28) groups compared to the negative control (7.31% ± 1.18, *p* < 0.05). Although the MeBB_250_ group (20.57% ± 4.67) also exhibited an increase in %OAT, this change was not statistically significant. The diazepam-treated group (33.14% ± 4.18, *p* < 0.05) demonstrated a significant increase compared to the negative control, and none of the MeBB-treated groups differed significantly from the diazepam-treated group. Additionally, the MeBB_250_ group differed significantly from the MeBB_1000_ group.

**Fig 3.**
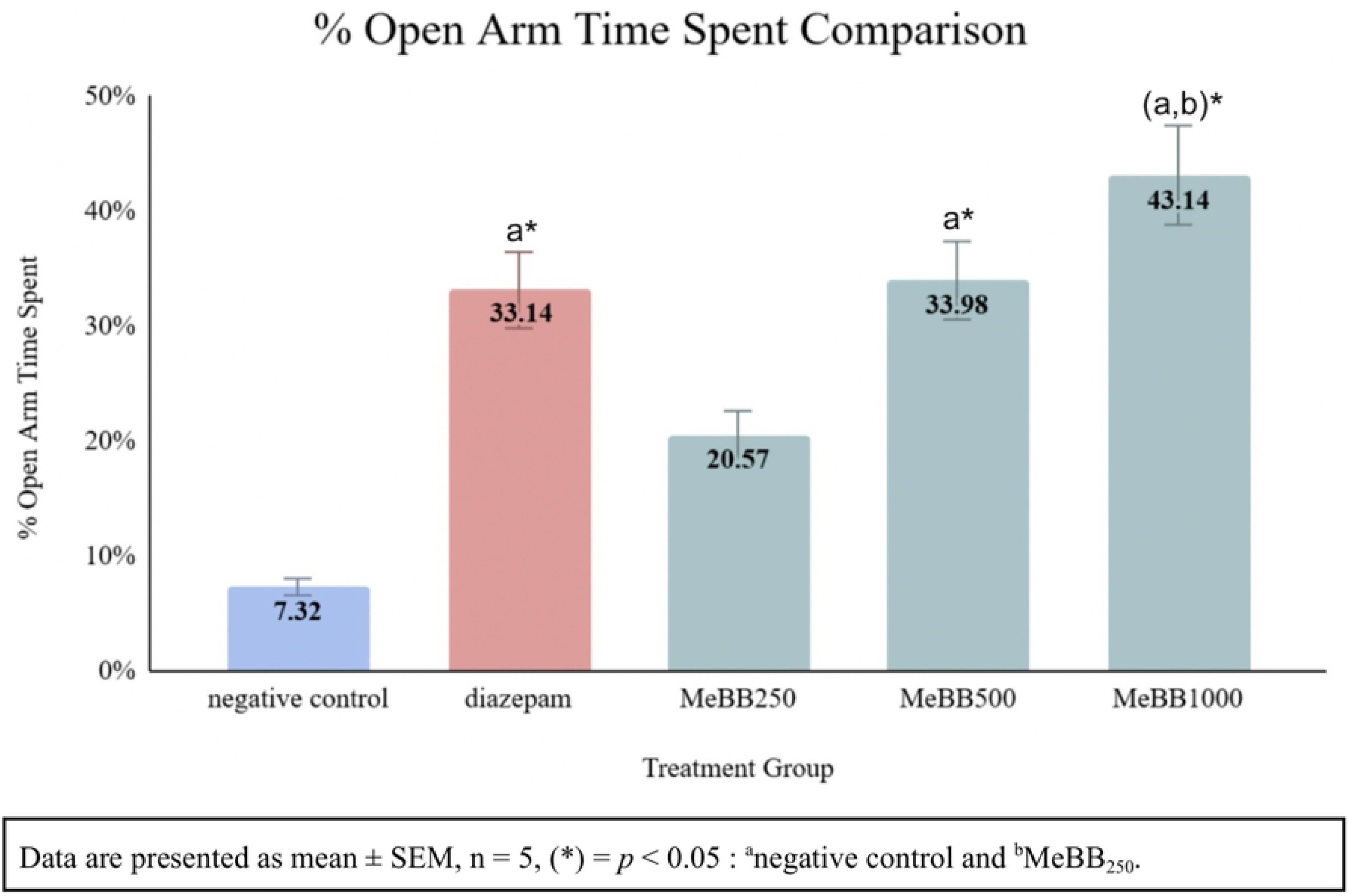
%OAT by the treatment groups in the EPM model.

A similar upward trend was observed in the percentage of open arm entries (%OAE), as illustrated in Fig 4. The MeBB_250_ (22.91% ± 5.19), MeBB_500_ (29.03% ± 4.25), and MeBB_1000_ (32.27% ± 9.80) groups showed numerical increases in %OAE compared to the negative control (18.52% ± 2.14, *p* < 0.05); however, these differences were not statistically significant. Likewise, none of the MeBB-treated groups differed significantly from the diazepam-treated group (34.92% ± 1.52, *p* < 0.05), and no significant difference was observed between the negative control and diazepam-treated groups.

**Fig 4.**
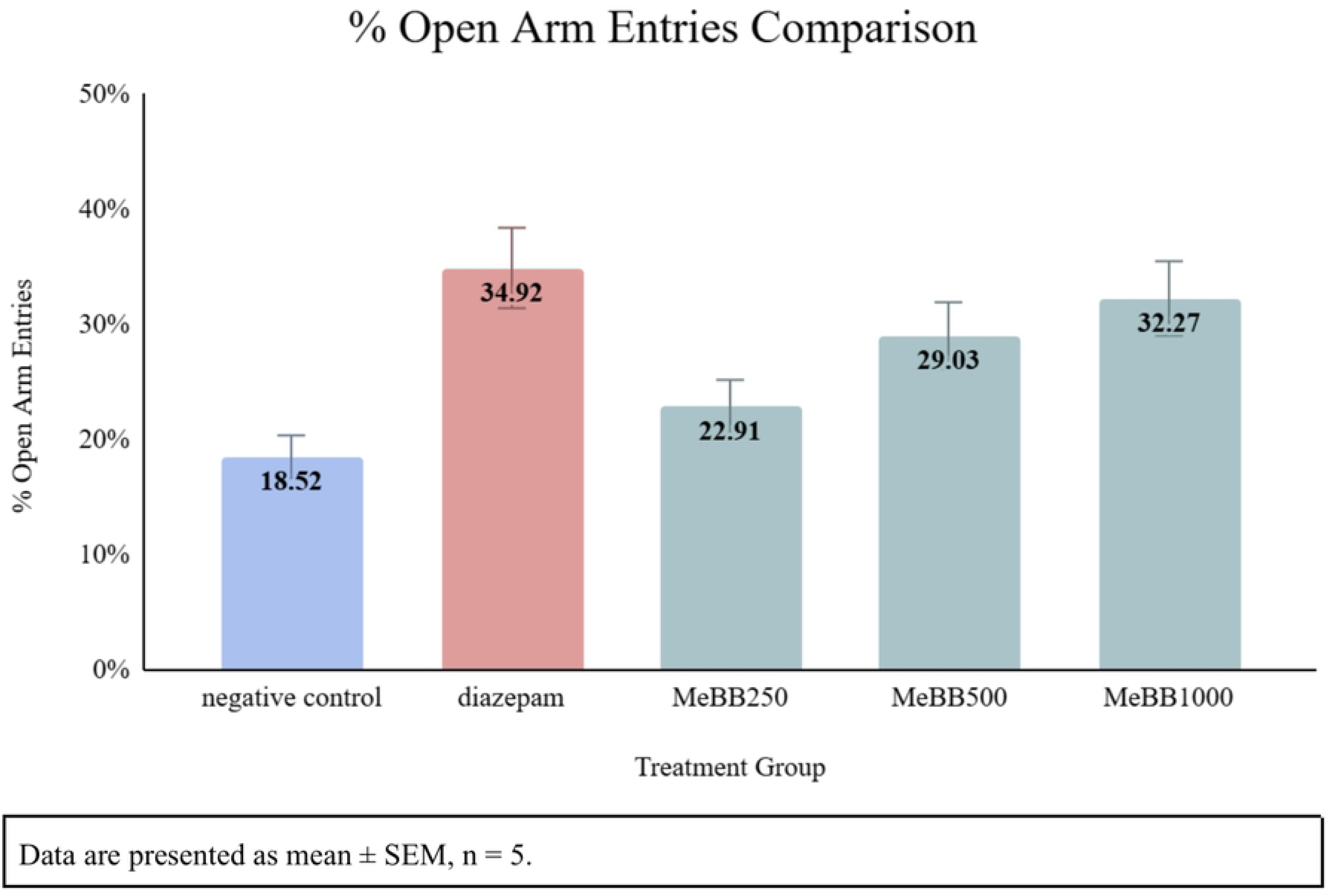
%OAE by the treatment groups in the EPM model.

#### Open arm time spent

This parameter measured the duration each mouse remained in the open arms of the maze. Mice naturally avoid open, elevated spaces due to their vulnerability to predators, so spending more time in these areas indicated reduced anxiety levels [7,8,55]. In this study, the observed increase in average open arm time among the MeBB-treated and diazepam groups, compared to the negative control, suggested a reduction in anxiety-like behavior.

#### Open arm entries

This parameter measured the number of times a mouse entered the open arms. A higher number of entries reflected reduced anxiety and increased exploratory behavior. However, this measure was also partly influenced by the mouse’s overall locomotor activity, making it necessary to interpret it alongside other activity-related parameters [8,13]. In this study, both open and closed arm entries showed an upward trend across all treatment groups, with the MeBB_1000_ group exhibiting the highest values in both. This pattern indicated that mice treated with the highest dose of the extract were not only more active but also more willing to enter the open arms, suggesting a combination of enhanced exploratory behavior and reduced anxiety-like responses.

#### Closed arm time spent

This parameter measured how long a mouse stayed in the closed arms. Since mice have a natural preference for enclosed areas, anxious individuals tended to spend more time in the closed arms while avoiding the open spaces [55]. In this study, a gradual decrease in average closed arm time was observed from the lowest to the highest MeBB dose, as well as in the diazepam-treated group, when compared to the negative control. This trend further supported the interpretation of reduced anxiety in the treatment groups.

#### Closed arm entries

This parameter measured how often the mouse entered the closed arms and was primarily used as an indicator of general locomotor activity rather than anxiety. It helped distinguish whether changes in open arm behavior were due to genuine anxiolytic effects or simply variations in movement, such as sedation or hyperactivity [13]. In this study, the similarity in trends between open and closed arm entries suggested that the extract, particularly at higher doses, might have produced a mild stimulant or arousal effect, increasing overall movement. This enhanced activity could have contributed to the increased number of open arm entries, not necessarily as a direct indicator of anxiolytic action alone. Therefore, interpreting open arm entries in the context of total locomotor activity was essential. The rise in closed arm entries helped confirm that the mice retained normal physical activity levels and were not sedated or behaviorally impaired, supporting the reliability of the observed anxiety-related behavioral changes.

#### Percentage of open arm time spent

This parameter measured the proportion of time the mouse spent in the open arms relative to the total time spent in both open and closed arms. A higher percentage of open arm time spent (%OAT) indicated reduced anxiety, as it suggested that the mouse was more willing to remain in exposed, elevated areas. Conversely, a lower %OAT reflected higher anxiety, as the mouse tended to avoid open spaces. Because %OAT accounted for individual differences in overall activity levels, it served as a more accurate and sensitive indicator of anxiolytic effects [56,57]. In this study, %OAT increased across all MeBB-treated groups and the diazepam-treated group, with the most pronounced effect observed at the 1000 mg/kg b.w. dose. Specifically, the MeBB_500_ and MeBB_1000_ groups showed statistical significance in %OAT values compared to the negative control, indicating a clear reduction in anxiety-like behavior. Although the MeBB_250_ group also showed an increase in %OAT, the difference was not statistically significant. The diazepam-treated group exhibited a %OAT comparable to that of the MeBB_500_ group. Importantly, none of the MeBB-treated groups differed significantly from the diazepam group, further supporting the potential anxiolytic effects of the extract at moderate to high doses.

#### Percentage of open arm entries

This parameter measured the proportion of total arm entries directed toward the open arms rather than the closed arms. A higher percentage of open arm entries (%OAE) reflected reduced anxiety and greater exploratory behavior toward exposed areas, while a lower %OAE suggested increased anxiety and preference for enclosed spaces. Like %OAT, this parameter adjusted for total locomotor activity and reflected not just movement, but the direction of exploratory behavior [13,55,56]. In this study, %OAE followed an upward trend across the MeBB-treated groups, but the differences were not statistically significant when compared to either control group. The similar upward trend of increase in both open and closed arm entries of the treatment groups suggested that the mice remained active and were not behaviorally suppressed, indicating that changes in anxiety-related behavior were not due to sedation or reduced movement. This general increase in movement likely influenced the total number of arm entries and may have balanced the ratio of open to total entries, minimizing differences in %OAE.

These refined behavioural parameters in the EPM (%OAT and %OAE) reflected two independent factors, anxiety-related behavior and general locomotor activity. Both are well-established indicators in anxiety research [13]. While some of the raw values for open arm time and entries did not show significant differences between the negative and positive control groups, the increase in %OAT observed in both the diazepam and higher-dosed MeBB groups indicated a clear reduction in anxiety-like behavior. The difference in statistical outcomes between %OAT and %OAE may be explained by increased overall activity, which affected the number of entries without necessarily altering the time spent in the arms. This made %OAE less sensitive in detecting treatment effects [57]. In addition, variations in individual locomotor activity likely contributed to inconsistent %OAE values. For example, some mice entered the open arms frequently but stayed only briefly, while others entered less frequently but remained longer. These behavioral differences were evident across treatment groups. In cases where total arm entries were low, small differences in the number of open arm entries could disproportionately impact %OAE values, further limiting its reliability. Overall, the consistent pattern of increased %OAT across treatment groups supported the conclusion that *B. balsamifera* extract, particularly at higher doses, reduced anxiety-like behavior, with %OAT emerging as the most reliable and consistent measure in this study.

The anxiolytic-like effects observed in the MeBB-treated groups were likely due to the combined activity of flavonoids, terpenoids, tannins, and phenols present in the crude methanol extract of *B. balsamifera*. These compounds had been reported to modulate neurotransmitter systems involved in anxiety regulation, particularly the GABAergic pathway, and may also have provided neuroprotective effects [35,36]. Their possible synergistic interaction may have contributed to the reduced anxiety-like behavior observed in the EPM. A similar finding was noted in *Cissampelos pareira*, whose extract contained various compounds such as alkaloids, flavonoids, tannins, and terpenoids and showed pharmacological effects likely due to synergistic interaction, although further investigation was required to confirm this [58].

Likewise, while the crude extract of *B. balsamifera* showed significant anxiolytic-like effects at 500 and 1000 mg/kg b.w., the specific contribution of each individual phytochemical remained unclear. The extract’s effectiveness only at high doses suggested that either the active compounds were present in low concentrations or their actions were diluted by the presence of inactive compounds. In contrast, diazepam, a purified and targeted drug, produced a similar behavioral effect at just 1 mg/kg b.w. Its high potency was due to its chemical purity and its well-established mechanism of directly enhancing GABA_A_ receptor activity, leading to rapid and reliable anxiolytic effects. According to Phootha *et al.* [36], plant extracts considered pharmacologically potent in animal models typically produce significant effects at doses below 30 mg/kg/day. Since *B. balsamifera* required much higher doses to demonstrate activity, it did not meet this potency standard. Consequently, these findings highlight a critical limitation regarding the immediate generalizability of the crude extract and emphasized the need to isolate and concentrate the active compounds in *B. balsamifera* to enhance its pharmacological strength, reduce the required dose, and better understand the mechanisms responsible for its anxiolytic-like activity.

## Conclusions

### Summary of findings

This study investigated the potential anxiolytic activity of the methanol extract of *Blumea balsamifera* (L.) DC. leaves through phytochemical screening, acute oral toxicity test, and an animal model of anxiety using the elevated plus maze (EPM) test. Mature leaves of *B. balsamifera* were collected from Guso, Jomgao, Argao Cebu, Philippines. These leaves were air-dried for 8-10 days, powdered, soaked in methanol for 48 hours, filtered, and concentrated using a rotary evaporator at 40°C. The resulting crude extract was screened for secondary metabolites and subjected to an acute oral toxicity test following OECD Guideline 423 to determine non-lethal concentration for behavioral testing. Behavioral responses were assessed using the EPM at doses of 250, 500, and 1000 mg/kg b.w., p.o., and were compared with a negative control (10 mL/kg b.w., p.o., 1% Tween 80) and a positive control (1 mg/kg b.w., p.o., diazepam).

Phytochemical screening of the crude extract identified naturally occurring compounds with reported central nervous system activity, particularly those associated with anxiolytic effects—namely flavonoids, terpenoids, tannins, and phenols. The acute oral toxicity test confirmed that the crude methanol extract of *B. balsamifera* leaves exhibited a relatively low toxicity profile, with no observed mortality or severe adverse effects at doses up to 2000 mg/kg b.w., p.o. Based on these findings, the crude extract was classified under Category 5 of the Globally Harmonized System, with an estimated LD₅₀ of 5000 mg/kg b.w. This justified the use of 250, 500, and 1000 mg/kg b.w. doses for subsequent behavioral testing. In the EPM test, mice treated with 500 and 1000 mg/kg b.w., p.o. of the crude extract showed a statistically significant increase in percentage of open arm time spent (%OAT) compared to the negative control, indicating anxiolytic-like activity. However, no statistically significant differences in the percentage of open arm entries (%OAE) were observed. These findings support a dose-dependent anxiolytic effect, particularly at higher doses, with %OAT emerging as the most reliable and consistent behavioral measure in this study.

### Concluding remarks

The results of this study suggest that the crude methanol extract of *Blumea balsamifera* (L.) DC. leaves possess anxiolytic activity. Thus, *B. balsamifera* shows potential as a natural anxiolytic agent, particularly at doses of 500 and 1000 mg/kg b.w.

### Recommendations

To fully explore the potential of *Blumea balsamifera* (L.) DC. as a natural anxiolytic agent, future research is strongly recommended to build on the current findings and further validate its therapeutic potential. It would be beneficial to repeat this study using other well-established animal models of anxiety, such as the light and dark box (LDB) and open field test (OFT), which could provide additional insights into the plant’s anxiolytic effects. Moreover, exploring the use of different solvents, including ethanol, n-hexane, acetone, and ethyl acetate, may yield extracts with varying metabolite profiles and activities, which could influence its effectiveness. Future research could also evaluate the anxiolytic activity of the volatile oils present in the plant.

Future studies should explore a wider range of doses and conduct comparative phytochemical analyses, including total flavonoid content and total phenolic content. These assessments will help clarify whether the observed anxiolytic activity at high doses was due to low concentration of active compounds or the presence of inactive compounds. Isolating and characterizing the bioactive constituents will be essential to improving potency, consistency, and understanding the underlying mechanisms of action.

This study contributes to the growing body of evidence suggesting that flavonoids, among other compounds, hold promise as leads in anxiolytic drug development. As the field continues to seek novel, safer treatments, *B. balsamifera* stands as a potential candidate for further exploration.

*“No formal study protocol was registered in a public repository prior to the initiation of the study. However, all experimental procedures, sample sizes, and analytical plans were reviewed and approved in advance by the University of San Carlos Institutional Animal Care and Use Committee (IACUC) under protocol number 2024–056–028.”*

*“All relevant data supporting the findings of this study are included within the manuscript and its Supporting Information files.”*

*“The authors have declared that no competing interests exist.”*

## Supporting information

S1 Appendix. Plant identification

S1 Fig. Plant identification certificate.
S2 Appendix. Phytochemical screening procedure and reagents
S3 Appendix. Certificate of IACUC approval

S2 Fig. Certificate of approval from USC IACUC.
S4 Appendix. Dosage stock solution concentration calculations

S1 Table. Preparation of stock solutions of the crude methanol extract of *Blumea balsamifera* (L.) DC. administered to mice used in the acute oral toxicity test.
S2 Table. Preparation of stock solutions of the crude methanol extract of *Blumea balsamifera* (L.) DC. administered to mice used in the elevated plus maze test.
S5 Appendix. Dose volume calculation for each mouse

S3 Table. Dose volumes administered per mouse in the acute oral toxicity test.
S4 Table. Dose volumes administered per mouse in the elevated plus maze test.
S6 Appendix. Body Scoring System (BCS)
S7. Appendix. Certificate of diazepam use for thesis purposes

S3 Fig. Certificate of authorization to use diazepam drug for research purposes.
S8 Appendix. Acute oral toxicity observation sheets
S9 Appendix. Gross necropsy data

S4 Fig. Gross necropsy performed on the mice following the acute oral toxicity test.
S5 Table. Gross necropsy of mice treated with 300 mg/kg b.w., p.o., of the crude methanol extract of *B. balsamifera* leaves.
S6 Table. Gross necropsy of mice treated with 2000 mg/kg b.w., p.o., of the crude methanol extract of *B. balsamifera* leaves.
S10 Appendix. % Moisture content data

S7 Table. % Moisture content of the coarse powdered *B. balsamifera* leaves.
S11 Appendix. Phytochemical screening data and observations S8 Table. Phytochemical screening data and observations.
S12 Appendix. Body weight gain
S9 Table. Calculations for body weight gain (%) from day of dosing to day 7 and 14.
S10 Table. Weekly body weights and body weight gain (%) from the day of dosing to day 7 and day 14 in mice treated with 300 mg/kg b.w., p.o., of the crude methanol extract of *B. balsamifera* leaves (Group 1, 1st step).
S11 Table. Weekly body weights and body weight gain (%) from the day of dosing to day 7 and day 14 in mice treated with 300 mg/kg b.w., p.o., of the crude methanol extract of *B. balsamifera* leaves (Group 2, 2nd step).
S12 Table. Weekly body weights and body weight gain (%) from the day of dosing to day 7 and day 14 in mice treated with 2000 mg/kg b.w., p.o., of the crude methanol extract of *B. balsamifera* leaves (Group 3, 1st step).
S13 Table. Weekly body weights and body weight gain (%) from the day of dosing to day 7 and day 14 in mice treated with 2000 mg/kg b.w., p.o., of the crude methanol extract of *B. balsamifera* leaves (Group 4, 2nd step).
S13 Appendix. Weights of liver, kidneys, and spleen of each mouse

S14 Table. Individual organ weights of mice treated with 300 mg/kg b.w., p.o., of the crude methanol extract of *B. balsamifera* leaves after a 14-day acute oral toxicity test.
S15 Table. Individual organ weights of mice treated with 2000 mg/kg b.w., p.o., of the crude methanol extract of *B. balsamifera* leaves after a 14-day acute oral toxicity test.
S14 Appendix. Photo documentation of elevated plus maze test

S5 Fig. (a) Top view of the elevated plus maze; (b) Area where mice were weighed prior to oral administration; (c) Setup surrounding the elevated plus maze test in the testing room.
S15 Appendix. Elevated plus maze test raw data

S16 Table. EPM data for mice treated with the negative control, 1% Tween 80, p.o.
S17 Table. EPM data for mice treated with the positive control, diazepam, p.o.
S18 Table. EPM data for mice treated with 250 mg/kg b.w. of *B. balsamifera* extract, p.o.S19
Table. EPM data for mice treated with 500 mg/kg b.w. of *B. balsamifera* extract, p.o.
S20 Table. EPM data for mice treated with 1000 mg/kg b.w. of *B. balsamifera* extract, p.o.
S16 Appendix. Statistical analysis data

S21 Table. One-way ANOVA results for the average open arm time spent in the EPM among treatment groups.
S22 Table. One-way ANOVA results for the open arm entries in the EPM among treatment groups.
S23 Table. One-way ANOVA results for the average closed arm time spent in the EPM among treatment groups.
S24 Table. One-way ANOVA results for the closed arm entries in the EPM among treatment groups.
S25 Table. One-way ANOVA results for the %OAT in the EPM among treatment groups.
S26 Table. One-way ANOVA results for the %OAE in the EPM among treatment groups.
S27 Tabe. Tukey’s HSD results for the average open arm time spent in the EPM among treatment groups.
S28 Table. Tukey’s HSD results for the open arm entries in the EPM among treatment groups.
S29 Table. Tukey’s HSD results for the average closed arm time spent in the EPM among treatment groups.
S30 Table. Tukey’s HSD results for the %OAT in the EPM among treatment groups.

